# Intra-abdominal resection of the umbilical vein and urachus of bovine fetuses using videolaparoscopy and celiotomy: surgical time and feasibility (cadaveric study)

**DOI:** 10.1101/2020.02.24.962431

**Authors:** Francisco Décio de Oliveira Monteiro, Heytor Jales Gurgel, Simon Silva de Sousa, João Pedro Monteiro Barroso, Gabrielle Patrizi Braga Vasconcelos, Daniele Lira dos Santos, Luiz Henrique Vilela Araújo, Loise Araújo de Sousa, Gabriela Melo Alves dos Santos, Kayan da Cunha Rossy, Verena Siqueira da Silva, Camila do Espirito Santo Fernandes, Barbara da Conceição Guilherme, Helaine Freitas Miranda, Carla Rozilene Guimarães Silva, Rodrigo dos Santos Albuquerque, Luisa Pucci Bueno Borges, Gilson Ferreira de Araújo, Pedro Paulo Maia Teixeira

**Author notes:** Corresponding author (FDOM). These authors contributed equally to this work (Heytor Jales Gurgel, Simon Silva de Sousa, João Pedro Monteiro Barroso, Gabrielle Patrizi Braga Vasconcelos, Daniele Lira dos Santos, Luiz Henrique Vilela Araújo, Loise Araújo de Sousa, Gabriela Melo Alves dos Santos, Verena Siqueira da Silva, Camila do Espirito Santo Fernandes, Barbara da Conceição Guilherme, Helaine Freitas Miranda, Carla Rozilene Guimarães Silva, Rodrigo dos Santos Albuquerque). These authors contributed equally to this work (Kayan da Cunha Rossy, Luisa Pucci Bueno Borges).

## Abstract

Surgical intervention for umbilical diseases in calves, when indicated, is a complementary and indispensable therapeutic resource for the treatment of umbilical conditions and is commonly performed using celiotomy. However, videolaparoscopy has demonstrated feasibility in many diagnostic and therapeutic procedures. The aim of this study was to assess the feasibility of the techniques and the surgical time of videolaparoscopy and celiotomy used in intra-abdominal resection of the umbilical vein and urachus of bovine fetuses (cadavers). Resection of the umbilical vein and urachus using videolaparoscopy and celiotomy was performed in 26 anatomical specimens (bovine fetuses obtained from an official slaughterhouse). Resection of umbilical structures was feasible with both techniques, but shorter surgical time and minimal tissue damage were achieved using videolaparoscopy. Laparoscopy requires specialized training and appropriate instruments and is an important tool for diagnostic and therapeutic exploration of the umbilical structures, liver, bladder, and associated/adjacent structures.

## Introduction

Umbilical disorders are often diagnosed in newborn bovine calves, and their etiologies may be associated with infections of the umbilical vessels and urachus, persistent urachus, and umbilical hernias [1]. In many cases, drug therapy alone is not sufficient to treat these conditions, and surgical intervention is a complementary and indispensable therapeutic resource for effective management of these disorders [2–4].

For the satisfactory and effective treatment of umbilical infections, reduction of bacterial load and removal of infected structures likely to have low antimicrobial penetration are indispensable as they significantly improve the prognosis of these diseases [5]. Surgical resection of umbilical structures is indicated for the treatment of calves with infected umbilical components and patent urachus and is usually performed using celiotomy [6–7].

Surgical resection of umbilical components using celiotomy has been shown to be effective for the treatment of omphalitis in calves, and it is easily performed by the surgeon using conventional surgical instruments, unlike laparoscopy, which requires special equipment and instruments that are usually more expensive than conventional ones, a fact that greatly influences their application in veterinary medicine [8–10]. In addition, performing laparoscopy requires the surgeon to have specific skills that can only be acquired with specialized training [11–13]. Therefore, the advantages of celiotomy make it the most used surgical technique in cases of umbilical conditions requiring surgical treatment [3,7].

Laparoscopy is a minimally invasive technique and an important alternative to celiotomy that allows, in addition to the observation of abdominal organs and structures, surgical techniques and exploratory procedures to be performed with less pain and better recovery for the patient than conventional techniques [13]. The use of laparoscopy in cattle is promising; there are many potential applications, and, because it is a safe and effective technique, it is an important alternative to conventional methods [11,14].

Surgical indication for the treatment of omphalitis should be preceded by a complete clinical examination, in particular an examination of the umbilical structures, with palpation of the umbilical region associated with ultrasound and/or laparoscopy, thereby allowing the surgeon to determine the severity of the abnormality and establish an appropriate surgical plan for the management of the specific case [10,15–17].

Data on the feasibility of the techniques and surgical time of procedures performed for anatomical specimens provide important information for the execution of these procedures in live animals. Thus, the objective of the present study was to assess the feasibility and the surgical time of videolaparoscopy and celiotomy for intra-abdominal resection of the umbilical vein and urachus of bovine fetuses (cadavers).

## Materials and methods

### Study site and groups

The experiment was conducted at the Institute of Veterinary Medicine (IMV) in Campus II of the Federal University of Pará (UFPA), located in the municipality of Castanhal/Pará, and involved the resection of the umbilical vein and urachus using videolaparoscopy and celiotomy in 26 anatomical specimens (bovine fetuses from cows in the final third of gestation slaughtered in an official slaughterhouse). The anatomical specimens were classified into two study groups: one group with 13 anatomical specimens that underwent resection of the umbilical vein and urachus using videolaparoscopy, i.e., videolaparoscopic surgery (GV, n = 13) and the other group with 13 anatomical specimens that underwent resection of the umbilical vein and urachus using celiotomy, i.e., open surgery (GA, n = 13).

All the steps of the procedures in both groups were performed by the same surgeon in a standardized and systematic manner throughout the study.

The feasibility of the techniques and the surgical time (both the total time and that of the steps of abdominal cavity entry, resection of the umbilical vein, resection of the urachus, and closure of the abdominal cavity) were analyzed in all the anatomical specimens in both groups. A descriptive comparison of both techniques was performed, in which the feasibility of performing the surgical procedures was assessed.

### Instruments and equipment used in the study

The experimental simulation of the surgical procedures took into consideration all the surgical principles applicable to videolaparoscopy and conventional open surgery, and the necessary equipment and instruments were used to perform the techniques. We used a 5-mm laparoscope, Babcock forceps, laparoscopic scissors, a set of gas insufflator/light source/monitor, and basic surgical instruments for conventional surgery.

### Access to the umbilical vein and urachus using videolaparoscopic surgery with three access ports

The anatomical specimens of the GV group were placed in the left lateral decubitus position and underwent videolaparoscopy using three videolaparoscopic access ports in the right flank, with two 10-mm cannulas in the first and second access ports and a 5-mm cannula in the third port for access to the umbilical vein and allantoic duct, or urachus. The access ports were established in the right flank near the paralumbar fossa, caudal to the ribs (Fig 1-A). Skin incisions of approximately 4 mm were made using a scalpel to insert the trocars transmurally inside the abdominal cavity, maintaining the triangulation of the access ports (Fig 1-B).

**Fig 1.**
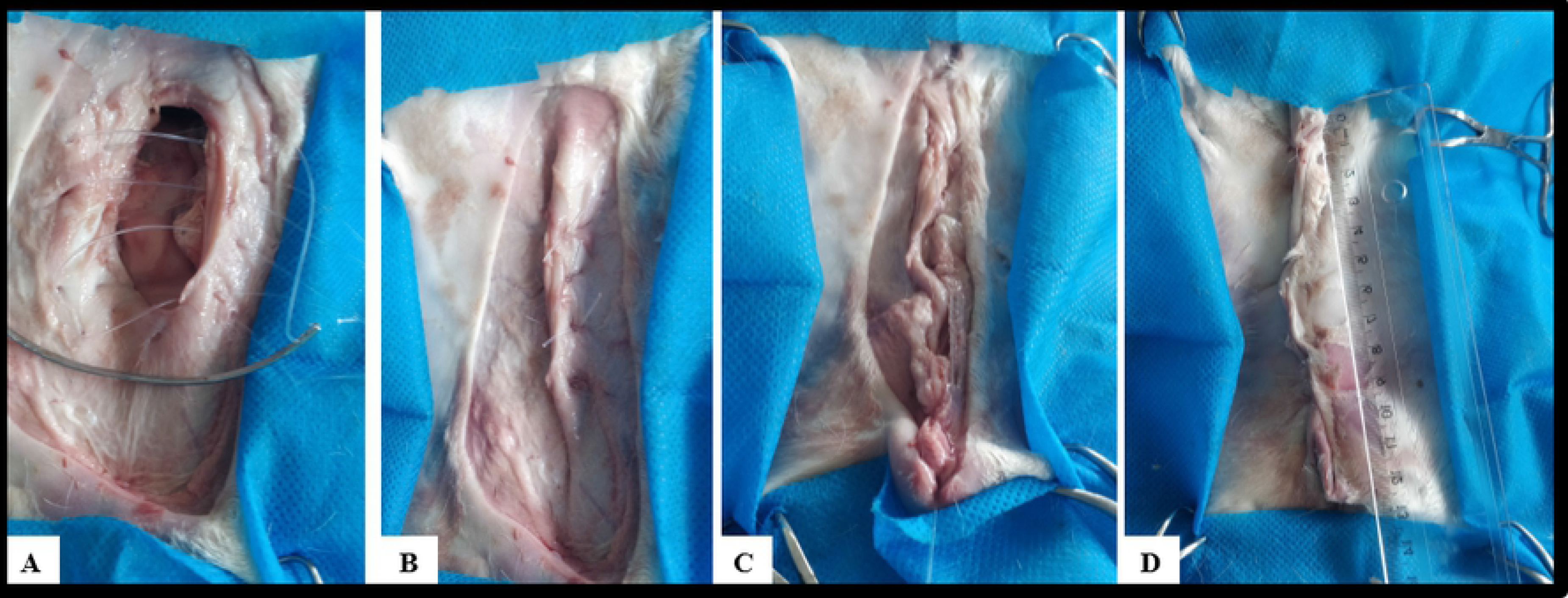
Positions of bovine fetuses that underwent videolaparoscopy and celiotomy for umbilical vein and urachal resection. Figs 1-A and 1-B show the flank region and triangulation of laparoscopic ports. Fig 1-C shows the umbilical and preputial regions and surgical field.

The first 10-mm videolaparoscopic access port with an insufflation valve was inserted, through which a carbon dioxide (CO_2_)-induced pneumoperitoneum of 8 mmHg was established, and the abdominal cavity was inspected by viewing the image on the monitor. The inspected elements included the umbilical vein from the umbilical ring to its insertion in the liver, the umbilical arteries from the umbilical ring to the vicinity of the bladder, and the urachus from the umbilical ring to the bladder.

The second videolaparoscopic access port was established using a 5-mm trocar to insert the Babcock forceps, through which manipulation of the umbilical structures was performed, simulating the clipping of these components and maintaining an appropriate position to perform the resections in the insertion regions and vicinity of the umbilical ring.

The third 5-mm port was established to perform the umbilical vein and urachal resection, in the insertion regions and near the umbilical ring, through which the laparoscopic scissors were introduced and resection of these components was performed. Subsequently, the components were removed through the second access port using the Babcock forceps.

### Resection of the umbilical vein using videolaparoscopic surgery with three access ports

For resection of the umbilical vein, the assistant operated the optics placed on the first port. The surgeon manipulated the umbilical vein, performed the clipping simulation, positioned the vein for resection, and resected it using laparoscopic scissors inserted in the third port. First, the surgeon placed the Babcock forceps in the second port and manipulated the umbilical vein for a better inspection of the regions to be resected, later removing the forceps. Subsequently, the Babcock forceps were reinserted to mimic a laparoscopic clipper used for hemostasis of large vessels, thereby simulating the application of four hemostatic clips in the umbilical vein: two in the vicinity of the umbilical ring and two in the vicinity of the liver, at the vein insertion site, and the Babcock forceps were again removed from the port. Finally, the Babcock forceps were reinserted to position and secure the umbilical vein. The surgeon placed the laparoscopic scissors through the third portal and resected the umbilical vein that was secured by the Babcock forceps at the “clipped” sites, and the vein was removed through the second access port using the Babcock forceps.

### Resection of the urachus using videolaparoscopic surgery with three access ports

The procedures for urachal resection followed the same sequence as those performed for umbilical vein resection described in the previous paragraph; however, separation of the tissues surrounding the urachus and bladder was performed using laparoscopic scissors introduced in the third port, for better identification of the “clipping” sites and resection. Simulation of clipping and urachal resection were performed near the umbilical ring and anterior to the bladder, following the same procedure used for simulation of clipping and resection of the umbilical vein.

### Closure of the abdominal cavity in videolaparoscopic surgery with three access ports

After resection of the urachus, the trocar cannulas and laparoscopic ports were removed, and the skin incisions were sutured with one stitch (U suture) using 0.6-mm polyamide thread.

### Access to the umbilical vein and urachus using celiotomy (conventional open surgery)

The anatomical specimens of the GA group were placed in the supine position with the hind legs abducted to better expose the umbilical and preputial regions, i.e., the surgical field (Fig 1-C). With the fetuses of bovine calves in the correct position and the surgical field established, a paramedian and circumscribed skin incision was made using a scalpel at the base of the umbilical region of each specimen. The subcutaneous tissue was pulled apart, and the muscles (rectus abdominis and external and internal oblique muscles) were incised down to the peritoneum. In the peritoneum, a stab incision was made using a scalpel, and the incision was enlarged using a scissors in the caudal and cranial direction and in a circumscribed manner, along the skin incision, to form a circumscribed opening in the abdominal cavity. This opening in the cavity allowed access to the umbilical structures, facilitating inspection and manipulation of the umbilical vein and urachus.

### Resection of the umbilical vein using celiotomy (conventional open surgery)

After access was established, the umbilical vein was inspected and a ligature was made with a Miller’s knot in the region adjacent to the liver, followed by two surgeon’s knots, and the vein was subsequently resected. Subsequently, the umbilical vein was removed from the abdominal cavity with an umbilical ring.

### Resection of the urachus using celiotomy (conventional open surgery)

After access was established, the tissues between the urachus and the artery were pulled apart for better inspection of the urachus and identification of the bladder, and two ligatures were made in the region near the bladder using Miller’s knots, followed by two surgeon’s knots for resection of the urachus, which was extracted from the cavity with umbilical ring.

### Closure of the abdominal cavity in celiotomy (conventional open surgery)

To seal the opening made in the abdominal cavity for celiotomy, peritoneum/muscle dieresis was performed with overlapping suture. The subcutaneous tissue was closed using simple continuous suture, and the dead space was reduced by anchoring the subcutaneous tissue to the musculature. The skin was closed with using U suture. All dieresis for closure of the abdominal cavity at this stage of the experiment was performed using 0.6-mm polyamide thread.

### Statistical analyses

The Shapiro–Wilk test was used to confirm that surgical time data were distributed normally. Data on total surgical time and duration of each surgical step were grouped for the GV and GA groups. Student’s t-test was used to compare normally distributed data on surgical time, and the Mann–Whitney test (Wilcoxon) was used for non-normally distributed data.

Descriptive statistics were processed using the R statistical program, version 3.0.2. The confidence interval was 0.95, and when p ≤ 0.05, the difference was considered significant and the null hypothesis was rejected.

## Results

All fetuses of bovine calves in the final third of gestation had well defined umbilical structures, including the components of interest, i.e., the umbilical vein and the allantoic duct, or urachus. These components had a cylindrical shape, defined walls, and flaccid texture, and it was possible to distinguish the umbilical veins and arteries as separate units. The liver and bladder were also well formed and defined. The bladder was empty, and the allantoic duct was cranially inserted in the anterior apex.

In the GV group, with the fetuses positioned in left lateral decubitus, it was possible to perform resection of the umbilical vein and urachus using videolaparoscopy with three access ports established in the right flank, in the paralumbar fossa region (Figs 1-A and 1-B). In the GA group, the most appropriate position for performing the proposed procedures was dorsal decubitus with abducted legs (Fig 1-C).

The pneumoperitoneum established using an intra-abdominal pressure of 8 mmHg allowed satisfactory separation of the abdominal wall from the visceral structures, as well as excellent intra-abdominal visualization of the umbilical structures and associated/adjacent organs such as the liver and bladder.

Access to the abdominal cavity and target structures to perform the procedures was achieved through three videolaparoscopic ports in videolaparoscopy and through a paramedian and circumscribed incision in the abdominal wall (subcutaneous tissue, muscle, and peritoneum) in celiotomy (Figs 1-B and 2). In videolaparoscopy, the surgical wounds consisted of three openings of approximately 3 to 5 mm in length, and in celiotomy the openings in the abdominal cavity ranged from 5.5 to 18.5 cm, with an average of 9.7 cm (Fig 2). Access to the abdominal cavity, down to the target structures, was secured in 5.52 (±1.82) and 4.78 (±0.74) min in the GV and GA groups, respectively (Table 1).

**Table 1.** Results of surgical time for each step of videolaparoscopy and celiotomy performed for resection of umbilical vein and urachus in bovine fetuses

| LEG | GV | GA | P Value |
| --- | --- | --- | --- |
| Access to the abdominal cavity | 5.52 ± 1.82 | 4.78 ± 0.74 | 0.1946 |
| Resection of the umbilical vein | 2.27 ± 1.33 | 2.89 ± 0.87 | 0.1734 |
| Resection of the urachus | 7.09 ± 1.27 | 3.13 ± 0.91 | <0.0001 |
| Closure of the abdominal cavity | 0.92 ± 0.09 | 28.64 ± 7.32 | <0.0001 |
| **Total Surgical Time** | **15.81 ± 3.26** | **39.44 ± 8.46** | **<0.0001** |

**Fig 2. Steps for intra-abdominal access to the umbilical vein and urachus in celiotomy.**

During access to the abdominal cavity using videolaparoscopy with three ports, the umbilical vein was the first component inspected because the laparoscope focused directly on the structure, thereby enabling its complete inspection (Fig 3-A). It was possible to simulate the clipping of the umbilical vein with Babcock forceps in the resection sites and extract it through the second access port using 10-mm Babcock forceps (Fig 3-A, 3-B and 3-C). Resection of the umbilical vein using celiotomy was achieved, and Miller’s knot was made to contain bleeding (Fig 4-B). The duration of umbilical vein resection was 2.27 (±1.33) and 2.89 (±0.87) min in the GV and GA groups, respectively (Table 1).

**Fig 3. Intra-abdominal resection of the umbilical vein and urachus of bovine fetuses using videolaparoscopy.**

Figs 8-A, 8-B and 8-C show the steps of umbilical vein resection. Figs 8-D and 8-E show the steps of urachal resection.

With regard to the urachus, it was possible to simulate clipping with the Babcock forceps and perform its resection using videolaparoscopy (Fig 3-E). Resection of the urachus using celiotomy was also achieved, and Miller’s knot in the duct was made in this case (Fig 4-C). The duration of urachal resection was 7.09 (±1.27) and 3.13 (±0.91) min in the GV and GA groups, respectively (Table 1).

**Fig 4. Intra-abdominal resection of the umbilical vein and urachus of bovine fetuses using celiotomy.**

In the GV group, closure of the abdominal cavity was performed by placing a stitch on the skin (U suture) that created a post-surgical wound of approximately 5 mm in length. In the GA group, peritoneum/muscle dieresis was performed with overlapping suture, subcutaneous tissue dieresis was performed with simple continuous suture, and dead space was reduced by anchoring the subcutaneous tissue to the musculature. Skin closure was performed with a U suture that created a post-surgical wound ranging from 10 to 20 cm, with an average of 13.6 cm (Fig 5). The surgical time for abdominal cavity closure was 0.92 (±0.09) min in the GV group and 28.64 (±7.32) min in the GA group (Table 1).

**Fig 5. Abdominal tissue and skin closure of the abdominal cavity.**

Figs 5-A and 5-B show peritoneum/muscle dieresis with overlapping suture. Fig 5-C shows reduction of subcutaneous dead space with muscle anchoring. Fig 5-D shows skin closure with U suture.

The total surgical time in the GV and GA groups was 15.81 (±3.26) and 39.44 (±8.46) min, respectively (Table 1).

## Discussion

The treatments indicated for umbilical vein and urachal diseases in newborn calves are based on clinical and/or surgical therapy, with conventional open surgery commonly performed in cases requiring surgical treatment. The aim of the present study was to assess the use of laparoscopy in umbilical vein and urachal interventions when surgical treatment is indicated. The surgical management of umbilical conditions usually has a good prognosis. The use of laparoscopy improves prognosis, with better postoperative results including less pain and better recovery for patients [3,18].

In GA, a satisfactory procedure involving the umbilical structures was possible, with adequate inspection and palpation of the umbilical structures, which allowed the surgeon to examine and resect each component. However, a more comprehensive exploration of the abdominal cavity was achieved using videolaparoscopy, with good intervention in the umbilical structures and adjacent tissues/organs because the visual field projected on the monitor enlarged the structures and improved the visualization of smaller areas. Using videolaparoscopy, intervention in tissues present in small areas and that are difficult to access is possible. This is a minimally invasive procedure that, when associated with advanced multimedia resources, enables new approaches and procedures that are more precise and objective, thereby improving the outcomes and reducing complications [11,14].

In the first step, there was no statistically significant difference in the time required to access the umbilical vein and urachus between the two groups. The duration of access in the GV group was longer because the time to insufflate the abdominal cavity was included in this step. In both groups, caution as well as the establishment of standardized access ports allowed the procedures to be completed without complications, both morphological complications and macroscopic lesions of abdominal structures. In videolaparoscopy, a skin incision smaller than the diameter of the trocar allowed the latter to be introduced slowly and carefully, thereby preventing potential injuries to the abdominal organs and CO_2_ losses from the cannulas of the laparoscopic ports during pneumoperitoneum. The trocars were inserted directly without injuring the abdominal organs and structures adjacent to the umbilical components and allowed working with instruments sufficiently far from the umbilical structures. Standardizing abdominal access techniques with safer and reliable methods reduces the likelihood of perioperative complications [14,19–20].

Although preventive measures were not adopted during CO_2_ insufflation in videolaparoscopy, it may be important to adopt them in live patients, especially in those with infected umbilical structures and with metabolic impairment, because pneumoperitoneum can induce hypothermia and respiratory and cardiovascular changes [2, 21–22].

The difference in the surgical times of umbilical vein resection (second step) between the GV and GA groups was not statistically significant. In videolaparoscopy, the time required for simulation of the clipping and umbilical vein resection was shorter than that for the execution of Miller’s knot followed by vein resection in conventional open surgery; however, the surgeon’s skill may have been a factor that influenced the surgical time of this step. A surgeon’s skill is directly related to specialized training that ensures the ability to perform procedures quickly and efficiently [11–13]. In videolaparoscopy, umbilical vein resection was performed with ease, and the positioning of laparoscopic ports, in particular, was very important to guarantee direct access to the structure, avoiding organs or tissues that could prevent or hinder the procedures and allowing resection to be performed safely and without complications [2,15].

The surgical time for urachal resection in videolaparoscopy was longer than that in conventional open surgery because access to the urachus in videolaparoscopy was not direct, similar to the access to the umbilical vein. The urachus lies in the ventral region of the abdomen, between the umbilical arteries, and is enveloped by tissues that make its identification difficult (not detected during laparoscopic visualization). For satisfactory intervention in the urachus, it is important to perform tissue divulsion around the duct, a procedure that requires a skilled laparoscopist and may have influenced the surgical time of this stage, although not greatly and not affecting the final surgical time. In general, urachal resection was achieved in all the animals in videolaparoscopy, which was shown to be an efficient technique for urachal interventions, both from a therapeutic and a diagnostic perspective. Considerable advances have already been achieved in the use of laparoscopy in urachal conditions [15,23].

In the final step, i.e., closure of the abdominal cavity, the surgical time in the GA group was longer than that in the GV group. This was expected because the opening of the abdominal wall in conventional open surgery was multiple times larger than that in videolaparoscopy, and additional time was required to perform abdominal tissue closure. In videolaparoscopy, a simple stitch ensured closure of the cavity, whereas in conventional open surgery, several stitches were required to perform peritoneum/muscle and subcutaneous dieresis and skin closure. Abdominal tissue closure in laparoscopy is satisfactorily performed with one or two simple stitches in each created wound and has lower rates of complications compared with abdominal tissue closure performed in conventional open surgeries [15,19].

The total surgical time of videolaparoscopy was significantly shorter than that of conventional open surgery (exactly 15.81 ± 3.26 min), which shows that resection using videosurgery of the umbilical vein and urachus is faster than resection using conventional open surgery. The step of abdominal cavity closure in conventional open surgery significantly affected the total surgical time in the GA group because access wounds in the abdominal wall were significantly larger than those caused by videolaparoscopy. This increased the time of abdominal cavity closure and, consequently, the total surgical time of conventional open surgery. The total surgical time of videolaparoscopy in the present study was significantly shorter than that in other similar studies in which the laparoscopy technique was used in diagnostic and surgical interventions of umbilical structures with times ranging from 36 to 160 min [2,15,24].

Abdominal wall injuries were less common in videolaparoscopy than in conventional open surgery; videolaparoscopy caused only three lesions of approximately 10 mm, as well as less tissue damage. This allows us to infer that videolaparoscopy causes less surgical trauma in living patients and provides better postoperative recovery of newborn calves. Because it is a minimally invasive technique, it is important to determine whether it is possible to perform it under regional anesthesia and sedation, a suitable alternative for treating calves with umbilical disorders associated with concomitant diseases and with reduced ability to adapt to general anesthesia [15]. Other studies have shown that laparoscopic techniques are less traumatic than conventional open techniques [25–28].

The results of videolaparoscopy allowed the identification of additional benefits of its use in production animals because the technique enabled a large visual field for the diagnostic exploration of the abdominal cavity, thereby providing diagnostic advantages from a macroscopic and morphological perspective, including the accurate detection of macroscopic disorders in the umbilical structures, liver, bladder, and other adjacent structures, with results that can be correlated with ultrasound and laparatomy. Videolaparoscopy is a potentially viable diagnostic alternative that allows diagnosing multifocal abscesses in the liver, changes in umbilical arteries near their branching from the internal iliac artery, and focal thickenings and adhesions [15,20,29]. The use of videolaparoscopy in production animals is still incipient, but its use provides better results in semiological and clinical/surgical procedures in bovine veterinary medicine, with advances in sanitary/production methods and animal welfare [18, 30–32].

## Conclusions

In both techniques, videolaparoscopy and celiotomy, resection of the umbilical vein and urachus was feasible; however, videolaparoscopy for adequate intra-abdominal procedures involving umbilical structures requires appropriate equipment and instruments in addition to specialized training.

The total surgical resection time of the umbilical vein and urachus using videolaparoscopy was significantly shorter compared with that using celiotomy. The longer duration of the latter was because of the large abdominal incision.

Both videolaparoscopy and celiotomy were feasible for resection procedures and for the inspection of umbilical structures and associated organs such as liver and bladder and can be used as important tools for diagnostic explorations.

## Approval by the ethics committee

CEUA/UFPA N° 6180180817

## Acknowledgements

We would like to thank the members of the VOR research team (UFPA) for their collaboration and contribution to this research work.

## References

1. Hopker A. Umbilical swellings in calves: a continuing challenge. Vet. Rec. 2014; 174: 219–220. doi: 10.1136/vr.g1790.

2. Bouré L, Foster RA, Palmer M, Hathway A. Use of an Endoscopic Suturing Device for Laparoscopic Resection of the Apex of the Bladder and Umbilical Structures in Normal Neonatal Calves. Vet Surg. 2001; 30: 319–326.

3. Williams HJ, Gillespie AV, Oultram JW, Cripps PJ, Holman AN. Outcome of surgical treatment for umbilical swellings in bovine youngstock. Vet Rec. 2014; 174(9): 221–221. doi: 10.1136/vr.101736.

4. Rodrigues CA, Santos PSP, Perri SHV, Teodoro PHM, Anhesini CR, Araújo MA, et al. Correlação entre os métodos de concepção, ocorrência e formas de tratamento das onfalopatias em bovinos: estudo retrospectivo. Pesqui Vet Bras. 2010; 30(8): 618–622. doi: 10.1590/S0100-736X2010000800002.

5. Reig Cordina L, Werre S R, Brown JA. Short-term outcome and risk factors for post-operative complications following umbilical resection in 82 Foals (2004-2016). Equine Vet J Suppl. 2018. doi: 10.1111/evj.13021.

6. Baird AN. Umbilical surgery in calves. Vet Clin North Am Food Anim Pract. 2008; 24(3): 467–477. doi: 10.1016/j.cvfa.2008.06.005.

7. Marchionatti E, Nichols S, Babkine M, Fecteau G, Francoz D, Lardé H, et al. Surgical Management of Omphalophlebitis and Long Term Outcome in Calves: 39 Cases (2008-2013). Vet Surg. 2016; 45: 194. doi: 10.1111/vsu.12433.

8. Mulon PY, Desrochers A. Surgical abdomen on the calf. Vet Clin North Am Food Anim Pract. 2005; 21(1): 101–132. doi: 10.1016/j.cvfa.2004.12.004.

9. Yanmaz LE, Okumus Z, Dogan E. Laparoscopic Surgery in Veterinary Medicine. Medwell J Vet Res. 2007; 1(1): 23–39. Available from: http://medwelljournals.com/abstract/?doi=vr.2007.23.39.

10. Nichols S. Umbilical surgery in calves. In: ProcMich Vet Conf. 2014. Available from: https://pdfs.semanticscholar.org/7a17/77e92de2ec1d10f423f0f080949d9a6948f5.pdf.

11. Buia A, Stockhausen F, Hanisch E. Laparoscopic surgery: A qualified systematic review. World J Methodol. 2015; 5(4): 238–254. doi: 10.5662/wjm.v5.i4.238.

12. Ferrufino FL, Cohen JV, Schaffner EB, Eulufi FC, Müller FP, Castillo JM, et al. Simulation in Laparoscopic Surgery. Cir Esp (English Edition). 2015; 93(1): 4–11. doi: 10.1016/j.cireng.2014.02.022.

13. Saber A, Bayumi EK, Sophia L, Hoek LSVD. Minimal Access and Minimally invasive Surgery in Veterinary Practice. J Surg. 2017; 5(3–1): 39–42. doi: 10.11648/j.js.s.2017050301.18.

14. Babkine M, Desrochers A. Laparoscopic surgery in adult cattle. Vet Clin North Am Food Anim Pract. 2005; 21(1): 251–279. doi: 10.1016/j.cvfa.2004.12.003.

15. Robert M, Touzot-Jourde G, Nikolayenkova-Topie O, Cesbron N, Fellah B, Tessier C, et al. Laparoscopic Evaluation of Umbilical Disorders in Calves. Vet Surg. 2016; 45(8): 1041–1048. doi: 10.1111/vsu.12559.

16. Seino CH, Bombardelli JA, Reis GA, dos Santos RB, Shecaira CL, Azevedo MR, et al. Avaliação ultrassonográfica de componentes umbilicais inflamados em bezerros da raça Holandesa com até 30 dias de vida. Pesqui Vet Bras 2016; 36(6): 492–502. doi: 10.1590/S0100-736X2016000600006.

17. Sato R, Yamada K, Shinozuka Y, Ochiai H, Onda K. Gas-filled urachal abcess with a pinging sound in a heifer calf. Vet Med. 2019; 64(8): 362–366. doi: 10.17221/61/2019-VETMED.

18. Patel AM, Parikh PV, Patil DB. Laparoscopy in veterinary practice. Vet Res Int. 2014; 2(1): 01–07.

19. Bianchi G, Martorana E, Ghaith A, Pirola GM, Rani M, Bove P, et al. Laparoscopic access overview: is there a safest entry method? Actas Urol Esp. 2016; 40(6): 386–392.

20. Oztoprak MY, Karaca I. Abdominal access techniques used in laparoscopic surgery. Int J Exp Clin Anat. 2019; 13(2): 129.

21. Rezende M, Prado O, Bandeira C, Petri A, Montero E. Body temperature evaluation during induced pneumoperitoneum with CO2: an experimental study in pigs. Surg Endosc. 2012; 26(6): 1724–1729. doi: 10.1007/s00464-011-2099-x.

22. Radlinsky MG. Complications and Conversion from Endoscopic to Open Surgery.Veterinary. Vet Clin North Am Food Anim Pract. 2016; 46 (1): 137–145. doi: 10.1016/j.cvsm.2015.07.004.

23. Marques LC, Marques JA, Marques ICS, Teixeira MCA. Dilatação cística do úraco e uroperitônio em touros: relato de cinco casos. Arq. Bras. Med. Vet. Zootec. 2010; 62 (6): 1320–1324. doi: 10.1590/S0102-09352010000600004.

24. Fisher AT Jr. Laparoscopically assisted resection of umbilicalstructures in foals. J Am Vet Med Assoc. 1999; 214(12): 1813–1816.

25. Easley JT, Garofolo SQ, Ruehlman D, Hackett ES. A 3-portal laparoscopic ovariectomy technique in ewes. Small Ruminant Res. 2014; 121(2–3): 336–339. doi: 10.1016/j.smallrumres.2014.08.001.

26. Zhang S, Hao M, Ma Y. Comparasion of laparoscopic and traditional abomasal cannulation in sheep. J Vet. 2016; 60: 113–117. doi: 10.1515/jvetres-2016-0016.

27. Tripathi SD, Khandekar GS, Datir AA, Ambore GS, Pawar A. Comparative Evaluation of Laparoscopic and Open Vasectomy Techniques in Teaser bulls. Intas Polivet. 2017; 18(2): 415.

28. Santos GMA, Barbosa AEC, Borges LPB, Morais HLM, Guilherme BC, Siqueira LS. Minimally invasive video-assisted ruminostomy in sheep. Small Ruminant Res. 2018; 167: 78–81. doi: 10.1016/j.smallrumres.2018.07.023.

29. Milovancev M, Townsend KL. Current concepts in minimally invasive surgery of the abdomen. Vet Clin North Am Small Anim Pract. 2015; 45(3): 507–22. doi: 10.1016/j.cvsm.2015.01.00.4.

30. Corrêa RR, Silva LCLC, Spagnolo JD, Castro LM, Andrade FSRM., Oliveira NFO. Evaluation of Ventral Laparoscopic Abomasopexy Using Surgical Staples Associated with Suture Material in Dairy Cattle. Acta Sci Vet. 2018; 46: 1545. doi: 10.22456/1679-9216.81834.

31. Gnus M, Ratajczak K, Antosik P. Application of a laparoscopic spieker of own design in reposition and fixation of the left displacement of the abomasum (LDA) in cattle. Med Weter. 2018; 74(2): 99–103. doi: 10.21521/mw.6059.

32. Sofi KA, Singh M, Kumar P, Sharma A. Laparoscopic chromopertubation for the evaluation of tubal patency in dairy cattle. Indian Journal of Animal Reproduction. 2018; 39(1).

